# Induced neural differentiation of human mesenchymal stem cells affects lipid metabolism pathways

**DOI:** 10.1101/2021.01.17.427010

**Authors:** Pnina Green, Inna Kan, Ronit Mesilati-Stahy, Nurit Argov-Argaman, Daniel Offen

## Abstract

Neuronal membranes contain exceptionally high concentrations of long-chain polyunsaturated fatty acids (PUFA), docosahexaenoic acid (DHA) and arachidonic acid (ARA), which are essential for neuronal development and function. Adult bone-marrow-derived mesenchymal stem cells (MSC) can be induced to possess some neuronal characteristics. Here we examined the effects of neuronal induction on the PUFA metabolism specific pathways. Differentiated cells contained ~30% less ARA than MSC. The expression of specific ARA metabolizing enzymes was upregulated, notably that of prostaglandin E_2_ synthase which increased more than 15-fold, concomitantly with a 3-fold increase in the concentration of PGE_2_ in the medium. Moreover, induced differentiation was associated with enhanced incorporation of exogenous DHA, upregulation of acyl-CoA synthases, fatty acid binding proteins, choline kinase (CK) and phosphatidylserine synthases as well as increased total cellular phospholipids (PL). These findings suggest that active ARA metabolites may be important in the differentiation process and that neuronal induction prepares the resulting cells for increased DHA incorporation through the action of specific enzymes.

## Introduction

One of the major features of neuronal tissue is the high proportion of the long-chain polyunsaturated fatty acids (PUFA) docosahexaenoic acid (DHA, 22:6 n-3) and arachidonic acid (ARA, 20:4 n-6) in neuronal membranes. These PUFA are important for multiple aspects of neuronal development and function, including neurite outgrowth [1–3], signal transduction, and membrane fluidity [4]. The incorporation of DHA in the nerve membranes during perinatal development is essential to the development of cerebral and retinal functions [5]. Moreover, DHA promotes the differentiation of neuronal stem cells in vitro and in vivo [6]. Although DHA is found in abundance in neuronal tissue, it cannot be synthesized by neurons and has to be supplied by cerebrovascular endothelium and astrocytes [7].

Long-chain PUFA are primarily incorporated into the *sn*-2 position of the phospholipids (PL). DHA is incorporated mainly into phosphatidylethanolamine (PtdEtn) and phosphatidylserine (PtdSer), whereas ARA is incorporated into phosphatidylinositol (PtdIns), PtdEtn and phosphatidylcholine (PtdCho) [8]. It has been repeatedly shown that DHA promotes PtdSer biosynthesis and accumulation [9,10] and inhibits apoptosis in neuronal cells in a PtdSer-dependent manner [10]. The increase of PtdSer levels upon DHA enrichment is not a universal mechanism, but specific to neuronal cells [11].

Adult autologous bone-marrow-derived mesenchymal stem cells (MSC) are able to differentiate beyond tissues of mesodermal origin into neuron-like cells, displaying a variety of neuronal attributesand markers of neuronal transmission [12]. Our group has previously reported a novel protocol for neuronal induction of MSC [13] and stressed the necessity of exogenous PUFA supplementation to this process [14]. Given the unique PUFA composition of neuronal tissue, we asked whether specific related metabolic pathways would be altered following induced differentiation. In the present study we report that, indeed, following neuronal induction specific changes occur, which include decreased ARA concentration, increased secretion of prostaglandin E_2_ (PGE_2_), increased incorporation of supplemented DHA, increased cellular PL, especially PtdCho and altered expression of related metabolizing enzymes. These results highlight the involvement of specific lipid pathways in neuronal development.

## Materials and methods

### Isolation and culture of human mesenchymal stem cells

Human MSC were purchased from Lonza, Inc. (Swiss) and cultured and expanded as previously described [15]. Briefly, cells were diluted 1:1 with Hank’s balanced salt solution (HBSS; Biological Industries, Beit-Haemek, Israel). Mononuclear cells were recovered from the fractionated suspension following centrifugation in Unisep-Maxi tubes (Novamed, Jerusalem, Israel) at 1000 g for 20 min at room temperature. The cells were resuspended in growth medium containing Dulbecco’s modified Eagle’s medium (DMEM; Biological Industries) supplemented with 15% fetal calf serum (FCS; Biological Industries), 2 mM L-glutamine (Biological Industries), 100 μg/mL streptomycin, 100 U/mL penicillin, 12.5 U/mL nystatin (SPN; Biological Industries) and plated in polystyrene plastic 75-cm2 tissue-culture flasks (Corning, Corning, NY). Non-adherent cells were removed after 24h with medium replacement. Adherent cells was cultured to 70%-90% confluence and reseeded at a density of 5,000-10,000 cells/cm^2^. The cells were maintained at 37°C in a humidified 5% CO_2_ incubator.

### Neuronal induction

Growth medium was replaced with Differentiation Medium I consisting of DMEM supplemented with 10% FCS, 2mM glutamine, SPN, 10 ng/mL basic fibroblast growth factor (bFGF; R&D Systems, Minneapolis, MN), 10 ng/mL epidermal growth factor (EGF; R&D systems), and N2 supplement (5μg/mL insulin; 20 nM progesterone; 100 μM putrescine; 30 nM selenium; 100 μg/mL transferrin). Seventy-two hours later, the medium was replaced with Differentiation Medium II containing DMEM supplemented with SPN, 2 mM L-glutamine, N2 supplement, 200 μM butylated hydroxyanisole (BHA; Sigma, St. Louis, MO), 1 mM dibutyryl cyclic AMP (dbcAMP; Sigma), 3-isobutyl-1-methyl-xanthine (IBMX; Sigma) and 1 μM all-trans-retinoic acid (RA; Sigma) for 48 hours.

### Serum deprivation

Growth medium was replaced with low serum medium containing DMEM supplemented with 10% FCS, 2mM glutamine and SPN. Forty-eight hours later the medium was changed into serum free medium containing DMEM, 2mM glutamine and SPN.

### Immunocytochemistry

Cells were fixed with 4% paraformaldehyde and permeabilized/blocked in 3% normal goat serum, 1% bovine serum albumin (Sigma), and 0.1 % Triton-X in PBS for 1 hour at room temperature. The cells were incubated with either mouse anti-β3-tubulin (TUJ1; 1:1000; Sigma) or mouse anti neuronal nuclei (NeuN; 1:10; Chemicon) for 24 hours at 4°C. NeuN staining was followed by goat anti-mouse Alexa-568 (1:1000, Molecular Probes). TUJ1 primary antibody was followed by goat anti mouse biotinilated (ready to use, Zymed) for 1 hour at room temperature followed by Streptavidin Alexa-488 (1:200, Molecular Probes). DNA-specific fluorescent dye 4,6-diamidino-2-phenylindole (DAPI; Sigma) counterstain was used to detect cell nuclei. Cells were photographed with a fluorescence Olympus IX70-S8F2 microscope with a fluorescent light source (excitation wavelength, 330–385 nm; barrier filter, 420 nm) and a U-MNU filter cube (Olympus, Center Valley, PA).

### Fatty acid supplementation and analysis

For the DHA incorporation experiments 30, 40, 50 or 60 μM DHA (Sigma) coupled with 1% horse serum (Biological Industries) and diluted in DMEM was supplemented to the growth medium or to the Differentiation Medium I.

For fatty acid analysis, following aspiration of the medium, cultures were washed with PBS, and lipids were extracted with hexane (BioLab, Jerusalem, Israel)/isopropanol (Sigma) (3:2, v/v) containing 5 mg/100 mL butylated hydroxytoluene (Sigma) as an antioxidant. Fatty acids were converted to fatty acid methyl esters (FAME) by heating with 14% boron trifluoride (BF3) in methanol (Sigma) and separated on capillary columns in an HP 5890 Series II GLC equipped with a flame ionization detector. Peak areas were integrated and plotted with the aid of the Varian Star Integrator computer package. Comparing retention times with authentic standards identified individual FAME. The amount of individual fatty acids is reported as percent peak area (area %) of total identified FA.

### Prostaglandin assay

PGE2 in the media of MSC and differentiated cells was determined by a PGE_2_ immunoassay (R&D Systems) according to the manufacturer’s instructions.

### Phospholipid analysis

Following aspiration of medium, cells were washed with PBS and stored at −20°C until lipid extraction. Total lipids were extracted from cells using a protocol adapted from the cold-extraction procedure developed by Folch et al. [16] with minor modifications, as previously described [17]. Briefly, total lipids were extracted from cells with methanol: chloroform: water solution (1:2:0.6, v/v). After overnight incubation at 4 °C, the lower, hydrophobic, phase was collected and filtered through a 0.45 μm Teflon syringe filter (Axiva Sachem Biotech, India) into a new vial. After solvents were evaporate under N_2_, total fat was dissolved in 100 μL of chloroform: methanol (3%, v/v) and stored at −20 °C until further analysis.

### Analysis of Polar and Neutral Lipids

Polar and neutral lipids were identified and quantified by HPLC (HP 1200, Agilent) combined with an evaporative light-scattering detector (ELSD, 1200 series, Agilent) according to a previously described protocol for normal-phase lipid separation [17]. Briefly, the separation protocol consisted of a gradient of dichloromethane, 99% methanol and 1% ammonium, and double-distilled water using normal-phase chromatography on a silica column (Zorbax RX-SIL, 4.6 × 250 mm, Agilent). The column was heated to 40°C, flow was 1 ml/min, and injection volume was 20 μl. Detector was heated to 65°C and nitrogen pressure was set on 3.8 bars. The separation process was managed by ChemStation software for data acquisition from the ELSD. Lipids were identified by external standards: PE, PS, PI, PC, and SM (Sigma Aldrich, Rehovot, Israel). Quantification was performed against external standard curves. Calibration curves were calculated by applying the power model equations to the concentration values, as following: PS, y = 0.0137X0.6339 (R^2^ = 0.999); PE, y = 0.0037X0.7398 (R^2^ = 0.987); PI, y =0.0135X0.5928 (R^2^= 0.999); PC, y = 0.0056X0.6661 (R^2^ = 0.99); SM, y =0.0101X0.6976 (R^2^ = 0.996).

### RNA extraction

Total RNA was isolated by a commercial Tryzol reagent (Invitrogen, Carlsbad, CA). The amount and quality of RNA was determined with the ND-1000 spectrophotometer (NanoDrop, Wilmington, DE) based on the ratios of 280/260 and 260/230. RNA was DNase (Qiagen, Germantown, MD) treated and cleaned using RNeasy Mini Kit (Qiagen). The overall quality of an RNA preparation was assessed by electrophoresis on a denaturing agarose gel.

### Microarray analysis

MSC derived from three human donors were included in the microarray study. Total RNA from untreated MSC and from differentiated cells was isolated as previously described. RNA labeling, chip hybridization, scanning and data acquisition were performed according to the manual provided by the Affymetrix manufacturer (http://www.affymetrix.com/support/downloads/manuals/wt_sensetarget_label_manual.pdf). Double-stranded cDNA was synthesized with random hexamers tagged with a T7 promoter sequence from 300 ng total RNA. cRNA was generated from the double-stranded cDNA template through an in-vitro transcription reaction and purified using the Affymetrix sample cleanup module. cDNA was regenerated through a random-primed reverse transcription using a dNTP mix containing dUTP. The RNA was hydrolyzed with RNaseH and the cDNA was purified. The cDNA was then fragmented by incubation with a mixture of UDG and APE1 restriction endonucleases and end-labeled via a terminal transferase reaction incorporating a biotinylated dideoxynucleotide. The efficiency of the labeling procedure was assessed by gel shift assay. 2.5 μg of the fragmented, biotinylated cDNA was added to a hybridization cocktail, loaded on a Human Gene 1.0 ST GeneChip and hybridized for 16 hours at 45°C and 60 rpm. Following hybridization, the array was washed and stained on a GeneChip Fluidics Station 450 according to the Affymetrix protocol. The stained array was scanned using an Affymetrix GeneChip Scanner 3000.

### Data analysis

Expression values from the Affymetrix Human Gene 1.0 ST chips were analyzed using Partek Genomics Suite (Partek Inc., St. Louis, MO). Raw intensity values were imported by setting up robust multiarray analysis background correction; quartile normalization, log transformation and median polish summarization. Principal components analysis (PCA) was performed for visualizing high-dimensional data. To identify differential expression between the groups, mixed-model ANOVA with linear contrast between two specific groups in the context of ANOVA, as implemented in the Partek software, was applied. *P*<0.01 cut off was applied and data was presented using geometric fold change.

### Quantitative reverse transcription polymerase chain reaction (real-time PCR)

Total RNA was isolated as previously described. First-strand cDNA synthesis was carried out with Super Script II RNase H-reverse transcriptase (Invitrogen, New Haven, CT) using a random primer. Real-time PCR was performed in an ABI Prism 7700 sequence detection system (Applied Biosystems, Foster City, CA) using Syber Green PCR Master Mix (Applied Biosystems) and the specific primers (Table 1). The glyceraldehyde 3-phosphate dehydrogenase (GAPDH) gene served as an internal control. The PCR was performed in a total volume of 20 μL containing 1 μL of the previously described cDNA, the 3’ and 5’ primers at a final concentration of 500 nM each, 7 μL of deionized H_2_O (water DEPC) and 10 μL of Syber Green Mix. The amplification protocol included 40 cycles of 95°C for 15 sec followed by 60°C for 30 sec. Quantitative calculations of the gene of interest versus GAPDH was performed according to the ΔΔCT method, as instructed in the user bulletin of the ABI Prism 7700 sequence detection system (updated 10/2001).

**Table 1:**
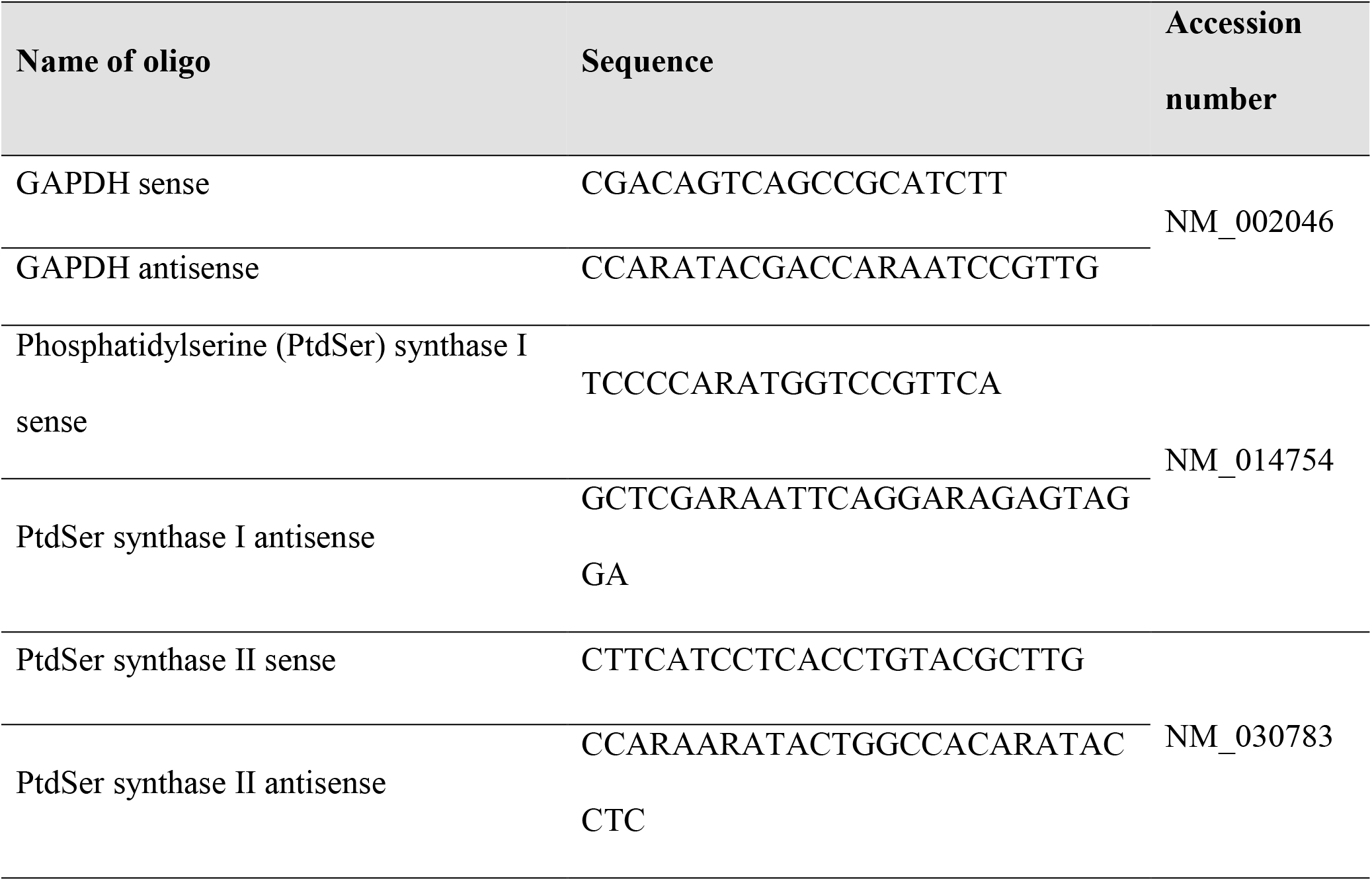
Oligonucleotide sequences of primers used for real-time PCR (Hylabs, Rehovot, Israel)

### Statistics

Data were analyzed using the SPSS software. Statistical analysis of two samples was performed by an Independent Samples two-tailed t-test. The statistical analysis of more than two samples was performed by ANOVA test (Scheffe post hoc comparison). The results were considered significant when *P*<0.05.

## Results

### Induced MSC show neuronal phenotype

Bright-light microscopy revealed that neuronal induction was associated with a morphological change of the cells, from the characteristic fibroblast shaped cells to a neuronal-like appearance (Fig. 1A, B). Immunocytochemical analysis revealed that the induced cells expressed higher levels of TUJ1, a neuronal progenitor marker, than the untreated MSC (Fig. 1E, F). In addition, the induced cells stained positive for the mature neuronal nuclear specific antigen NeuN (Fig. 1C, D).

**Figure 1:**
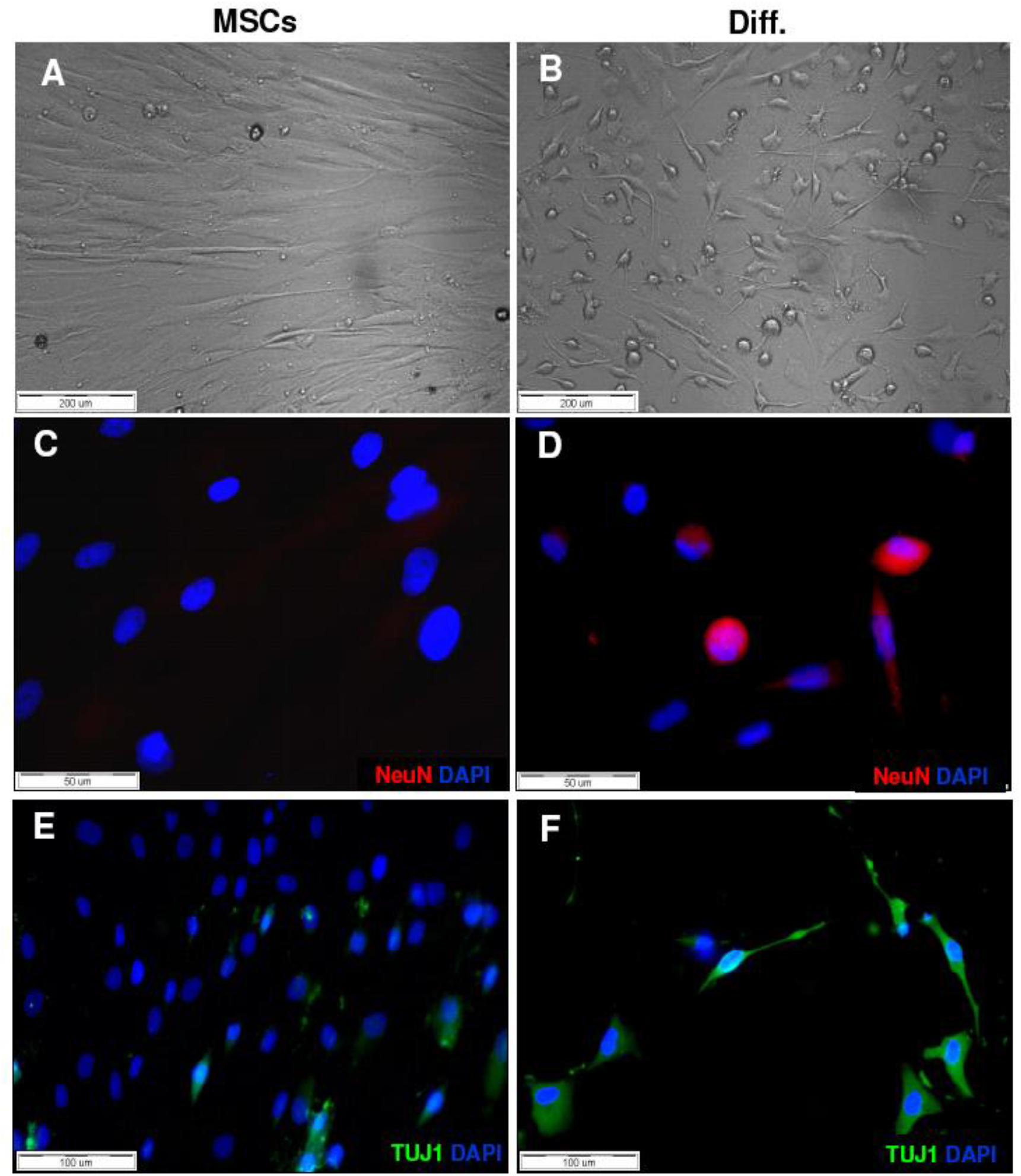
Neural characteristics of MSC following the induction. (A) Fibroblast-like morphology of MSC. (B) Neural morphology of MSC following the induction (C-F). Immunocytochemistry for neuronal markers TUJ1 and NeuN in induced cells (Diff.) and untreated MSC (MSC).

### Neuronal induction reduces the cellular concentrations of ARA

Following neuronal induction, a consistent decrease of ARA concentration was observed, from 8.37 ± 0.51 area % to 6.1 ± 0.64 area % (*P*=0.03, 22 samples of seven independent experiments) (Fig. 2). The only other significant change was an increase in the concentration of 20:3n-6 (from 1.57 ± 0.12 area % to 2.15 ± 0.15 area %). No differences were found in the n-3 PUFA or the other fatty acid subgroups (not shown).

**Figure 2:**
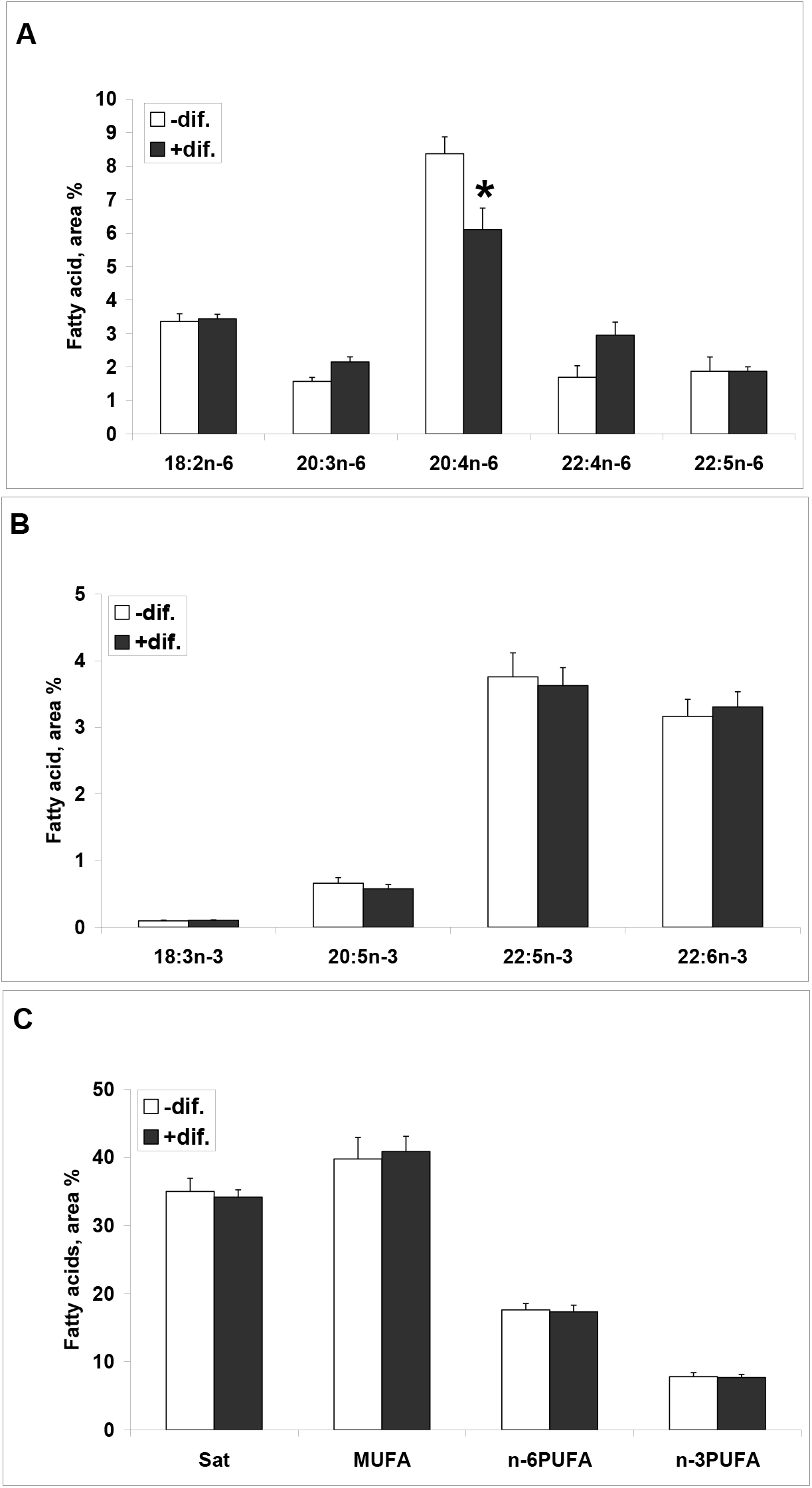
Fatty acid concentration of MSC upon neural induction. The fatty acid composition of the MSC and differentiated cells (Diff.) was determined by GLC of fatty acid methyl esters, as described in Materials and Methods (n=7 independent experiments). Values are expressed as area percentage of the total identified fatty acids, means + SE. *p<0.05

### The decreased concentration of ARA is not due to non-specific stress

Since neuronal induction involves removal of serum from the culture upon the final step of culture in Differentiation Medium II (Materials and Methods), a procedure which entails cellular stress [18], we checked whether serum removal might cause the decreased ARA. In MSCs without differentiation the concentration of ARA was 10.12 ± 1.07 area%, whereas upon serum removal the value remained virtually unchanged: 10.65 ± 0.17 area%. Thus, serum removal did not affect the cellular concentration of ARA.

### Specific pathways of ARA metabolism are affected by neuronal induction

Gene expression analysis related to ARA metabolism was performed in order to elucidate the observed decrease of ARA concentration. Microarray analysis of mRNA expression of MSC obtained from three human donors, before and after neuronal induction, included significant changes in phospholipases, in the cyclooxygenase pathway and the cytochrome P450 pathway (Fig. 3 and Supplements Table 1–4). Figure 3 is a cartoon depicting the major metabolic pathways of ARA in which the observed changes are noted. In particular, the phospholipases A_2_ (PLA_2_) increased 2-4 fold, cyclooxygenase 2 (COX_2_) increased almost 42-fold and prostaglandin E_2_ (PGE_2_) synthase increased 15-fold. Downregulation of prostacyclin (PGI_2_) synthase more than 9-fold was also noticed. No significant changes were recorded in the major enzymes of the lipoxygenase pathway and only a minor downregulation was observed in the cytochrome P450 pathway (Fig. 3).

**Figure 3.**
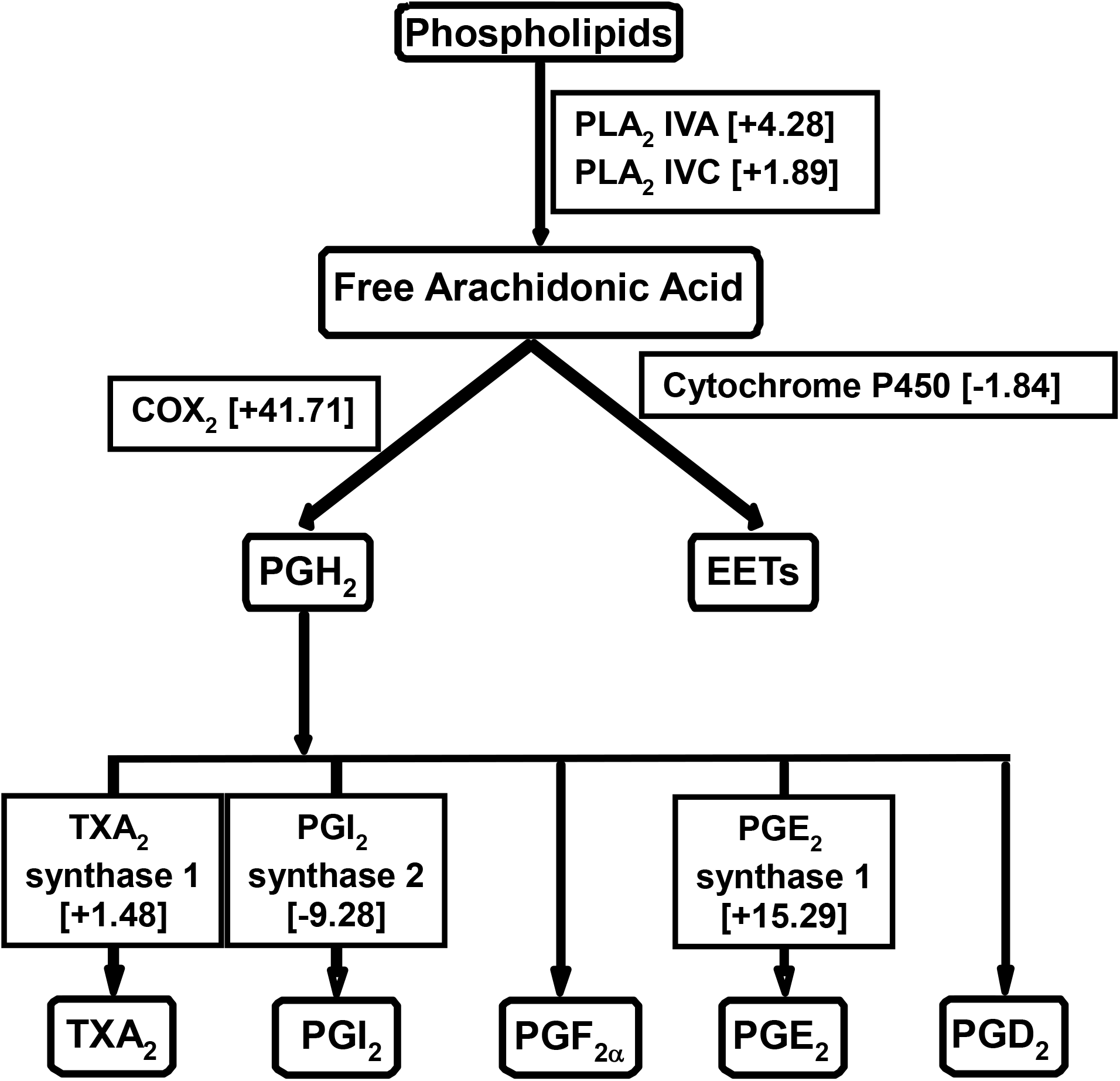
Pathways of arachidonic acid metabolism affected by neural induction. MSC were obtained from three human donors and induced to differentiate into neural cells. The change in mRNA levels of ARA metabolizing enzymes that resulted from the induction was evaluated by gene microarray as specified in Materials and Methods. The specified data represent enzymes that recorded significant change (p<0.01) following the neural induction and their fold change (positive values stand for upregulation and negative values represent downregulation). PG, prostaglandin; TX, thromboxane. PLA, Phospholipase; COX, Cyclooxygenase; EET, epoxy-eicosatrienoic acids.

### Neuronal induction is associated with increased PGE_2_ secretion

Immunoassay of the media for PGE_2_ from undifferentiated and differentiated cells showed a 3-fold increase upon differentiation induction (Fig. 4), in accord with the major upregulation observed for the enzymes participating in its synthesis.

**Figure 4.**
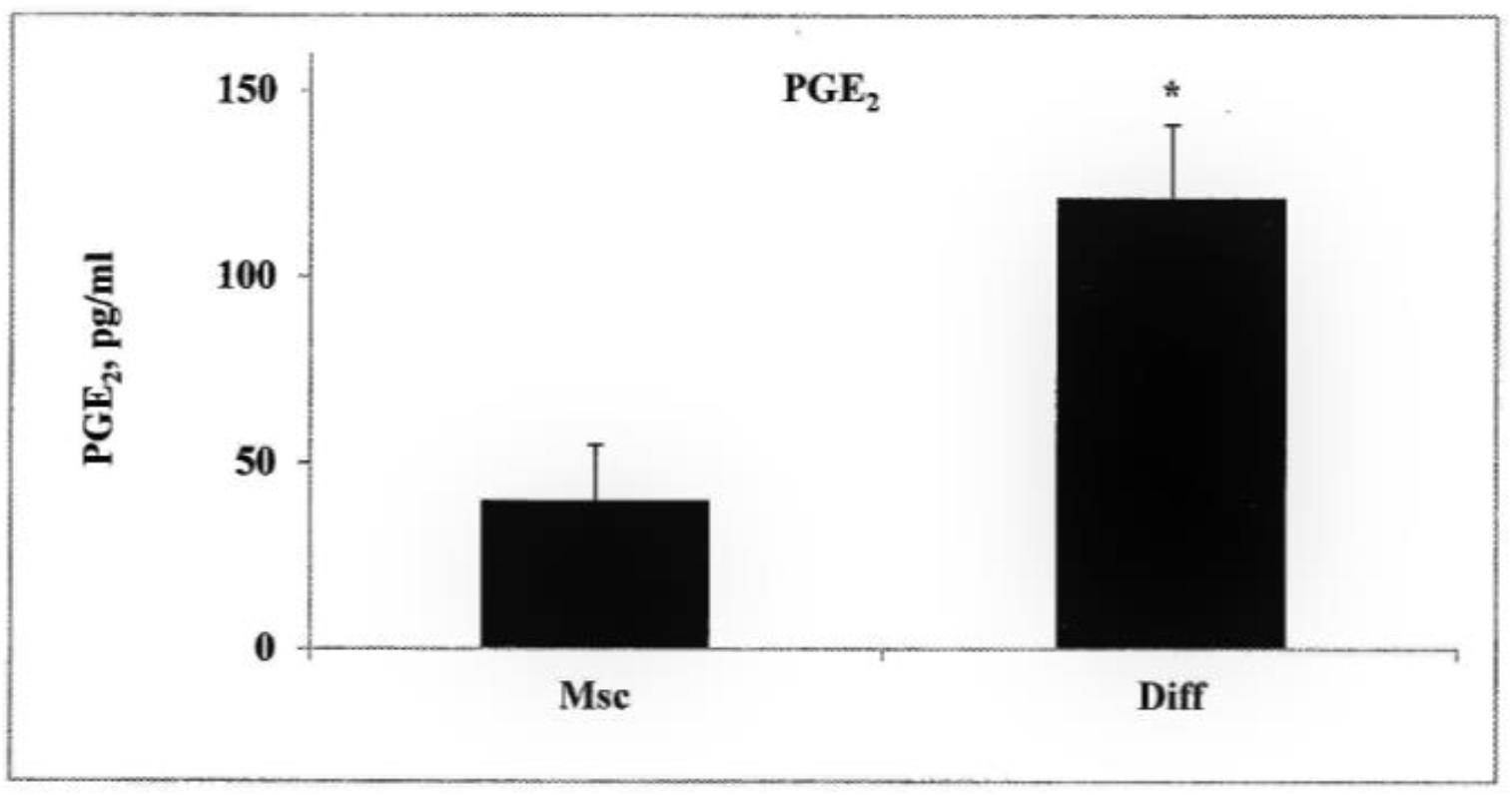
Prostaglandin E_2_ secretion upon neural induction. PGE_2_ in the media of MSC and differentiated cells (Diff) was determined by a PGE_2_ immunoassay. Values are mean + SE, n=5, p<0.03.

### Neuronal induction affects the incorporation of supplemented DHA

MSC following neuronal induction (Diff.) demonstrated considerable incorporation of DHA as opposed to undifferentiated MSC (Fig. 5). In dose-response experiments, differentiated cells incorporated more DHA at various supplemented concentrations, whereas the corresponding curve in the MSC remained almost flat. Specifically, the DHA concentration in differentiated cells supplemented with 30 μM DHA was 11.58 ± 1.92 area % vs. 7.87 ± 0.82 area % in undifferentiated cells supplemented with identical concentration of DHA. Following the supplementation of 40 μM DHA, DHA concentration was 15.88 ± 0.83 area % in differentiated cells vs. 8.36 ± 0.12 area % in undifferentiated cells. While the cellular concentrations of DHA kept climbing in the differentiated cells with increasing concentrations of the supplemented DHA, to 17.54 ± 1.19 area % following supplementation of 50 μM DHA and to 21.60 ± 0.21 area % following supplementation of 60 μM DHA, the cellular concentrations of DHA in the undifferentiated cells remained unchanged (~8.5 area %).

**Figure 5.**
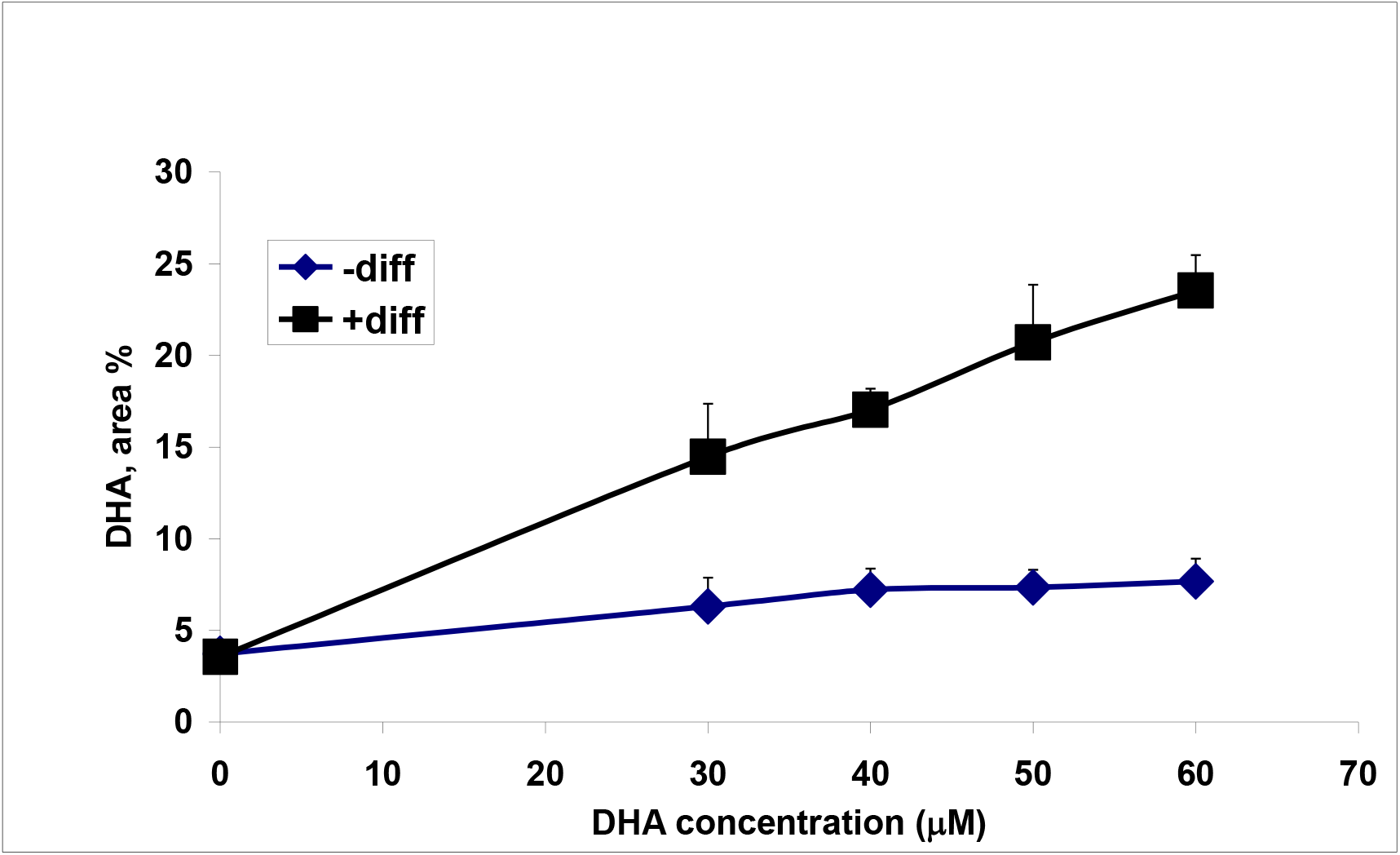
Dose response analysis of DHA supplementation. The concentration of DHA in MSC and differentiated cells (Diff.) cultured in the presence of various concentrations of DHA was determined by GLC, as described in Materials and Methods (two independent experiments, n=4 in each experiment). Values are expressed as DHA area percentage of the total identified fatty acids, means ± SE.

In the differentiated cells, an approximately 45% decrease was observed in the total monounsaturated fatty acids (MUFA) upon increasing the added DHA from 0 μM to 60 μM and the total n-6 PUFA decreased by approximately 18%. The saturated fatty acids remained unchanged (not shown).

### Microarray analysis of mRNA expression of enzymes and proteins involved in PL and PUFA metabolism is affected by neuronal induction

The main changes in PL synthesizing enzymes before and after the neuronal induction involved upregulation of PtdCho and PtdSer synthesis, namely, that of choline kinase (CK) which increased more than two-fold and of PtdSer synthases I and II, both which increased 1.5-fold (Data not shwen).

Enzymes involved in PUFA metabolism, which were affected by neuronal induction included a more than 10-fold upregulation of fatty acid binding protein 5 (FABP5), almost 6-fold upregulation of acyl-CoA synthetase long-chain family member 1 (ACSL1), almost 3-fold upregulation of ACSL3 and more than 11-fold upregulation of ACSL4 (Table 2 and Supplements Table 5).

**Table 2:** The fold change in the expression of fatty acid binding proteins and Acyl-CoA synthetase long-chain family members that occurred upon the induction and was found significant (p<0.01) in the microarray analysis.

| **Enzyme** | **Accession**<br>**number** | **P (+Diff. vs -**<br>**Diff.)** | **Fold Change** |
| --- | --- | --- | --- |
| FABP 5 | NM_001444 | 7.29E-06 | 10.24 |
| Acyl-CoA synthetase long-chain<br>family member (ACSL) 1 | NM_001995 | 6.11E-07 | 5.77 |
| ACSL 3 | NM_004457 | 0.001 | 2.95 |
| ACSL 4 | NM_022977 | 6.6E-05 | 11.43 |

### Real-time PCR

Analysis of PL synthesizing enzymes showed that neuronal induction increased the expression of CTP:phosphocholine cytydyltransferase beta (CCTb), PtdSer synthase I and PtdSer synthase II, enzymes which participate in the synthesis of PtdCho and PtdSer (Fig. 6).

### Cellular PL composition is affected by neuronal induction

Total cellular PLs increased in the cells following neuronal induction (74.7±9.4 mg/10^6^ cells vs. 43.1±8.0 mg/10^6^ cells, p=0.01), mainly due to increase of PtdCho (45.2±5.7 vs. 22.9±3.8mg/10^6^ cells, p=0.005).

## Discussion

In this study we demonstrated several changes involving lipid metabolism in MSC undergoing neuronal induction, especially in the cellular content of ARA and DHA. The findings related to ARA may point towards possible role of its metabolites in neuronal differentiation, whereas those related to DHA suggest that cells differentiating into neural cells are “primed” to increase their DHA accretion.

One of the effects of the neuronal induction was the decrease in the cellular levels of ARA. This decrease was induced by defined conditions of the induction protocol and did not occur following unspecific stress caused by serum deprivation.

The observed upregulation in the expression of PLA_2_ (more than 4-fold) and PGE_2_ synthase 1 (more than 15-fold), together with a 3-fold increase in PGE_2_ secretion are probably related to the decreased ARA, as detailed below.

Enzymatic oxidation of ARA includes three major metabolic pathways: COX, lipoxygenases (LOX) and epoxygenases (EPOX) which metabolize ARA to prostaglandins (PG), thromboxanes, leukotrienes, and epoxyeicosatrienoic acid, respectively. These metabolites are collectively referred to as eicosanoids and play important roles in regulating signal transduction and gene transcription processes [19]. Although the function of PG in the CNS has been studied mostly in relation to disease, eicosanoids in general and PG in particular, have important physiological functions. They play important roles in neural function including sleep induction, long term potentiation, spatial learning and synaptic plasticity [20]. Specifically, the stimulation of pheochromocytoma PC_12_ cells with nerve growth factor (NGF) results in outgrowth of neurites. This stimulation increases arachidonate metabolism and eicosanoid production, while inhibitors of PLA_2_ and lipoxygenase metabolism prevent neurite outgrowth [21]. Moreover, retinoic acid, a component of the differentiation medium in the present study and a well-known mediator of differentiation in neural cultures, has been postulated to be involved with the release of ARA by PLA_2_ and generation of eicosanoids [22]. From these considerations we suggest that the observed ARA decrease might be essential for the differentiation process, by the generation of oxidized active ARA metabolites.

The fundamental difference between MSC and differentiated cells regarding DHA accumulation in the present study is shown in Figure 5. It is well known that neuronal tissue is especially enriched in DHA, and here we demonstrate the profound change that cells undergo once they are induced to differentiate toward neuronal cells, leading to their ability to avidly accumulate supplemented DHA. In order to unravel some of the fatty acid-related mechanisms leading to neuronal differentiation, including the ability of neuronal tissue to accumulate DHA, we compared the expression of several related enzymes and proteins in MSC before and after neuronal induction (Fig. 3, 6, Table 2, Supplements). Specifically, a significant upregulation was recorded in the levels of the fatty acid binding protein (FABP) 5 (EFAB), of the acyl-CoA synthases ACSL1, ACSL 3 and ACSL 4, and a significant increase in the expression of choline kinase, PtdSer synthase I and PtdSer synthase II.

While it remains controversial how fatty acids cross the plasma membrane, considerable evidence implicate diffusion through the PL bilayer [23] as well as the involvement of several proteins – FABP - as facilitators of free fatty acid transport across the lipid bilayer [24]. FABP belong to the family of the intracellular lipid-binding proteins that are expressed in vertebrate tissues. FABP enhance both fatty acid uptake and intracellular esterification, but distinct FABP types differentially affect these processes [25]. Expression of three FABP types was detected in the brain: FABP3, FABP5 and FABP7 [24,25]. Studies with PC_12_ cells revealed that FABP5 (formerly known as EFABP) has high affinity to a broad range of fatty acids involved in axonal growth and is needed for normal neurite formation during neuronal differentiation [26]. In a mouse model it was demonstrated that FABP7 and FABP5 play essential roles in postnatal differentiation of neural stem cells [27], while in another study it was shown in vitro and in vivo that FABP5 binds to DHA and is involved in the brain endothelial cell uptake and subsequent supply of this essential fatty acid to the brain [28]. In our study, we found a significant upregulation of FABP5 (EFABP) upon neuronal induction of MSC, whereas FABP3 (HFABP) was slightly down-regulated and the expression of FABP7 (BFABP) did not change. (Supplement Table 6). We suggest that the increased expression of FABP5 following differentiation induction could be associated with the impressive accumulation of DHA in the differentiated cells (Fig. 5).

Following internalization, fatty acids are activated primarily by ACSL. Five mammalian ACSL have been identified. ACSL1 and ACSL5 activate most unsaturated FA similarly, whereas ACSL3, ACSL4, and ACSL6 (formerly ACS2) preferentially activate PUFA [29]. We observed the upregulation of three of the ACSL following neuronal differentiation (Table 2). Particularly noteworthy is the 11.4-fold increase in the expression of ACSL4, which has been shown to have a very high affinity for both ARA and DHA [30] Moreover, it is highly expressed in brain and is involved in neural differentiation through its role in ARA metabolism [31]. Therefore, this upregulation is consistent with the observed changes in both fatty acids following neuronal induction, namely decreased ARA and increased incorporation of DHA. Interestingly, the expression of ACSL6 remained unchanged (Supplement Table 5).

Neuronal differentiation is associated with increased neurite outgrowth and membrane synthesis, which involve increased PtdCho synthesis [32]. In the present study, the almost 2-fold increase in the PtdCho levels in the differentiated cells was associated with a more than 2-fold increase in the expression of CK, the first enzyme of PtdCho synthesis [32], as well as a significant increase in the expression of CCTb (Fig. 6).

Upon differentiation induction we observed upregulation of the expression of PtdSer synthase I and PtdSer synthase II, which catalyze the synthesis of PtdSer from PtdCho and PtdEtn, respectively (Fig. 6). The amount of PtdSer remained unchanged following differentiation, but it is expected to increase with DHA supplementation (Reviewed in [10]).

In summary, we show that neuronal induction of MSCs results in changes of lipid elements characteristic of neuronal cells. Further, we demonstrate some of the mechanisms involved in the PUFA metabolic pathways following neuronal induction. We suggest that active oxidized metabolites of ARA, especially PGE_2_, may be important for the differentiation process. Specific enzymatic pathways are induced to facilitate the incorporation of cellular DHA and the synthesis of PtdSer: the latter’s importance together with DHA for proper neuronal function and survival has amply been documented. Extended lipidomic analysis and detailed examination of involved enzymes should be the supplementary steps in future studies. We hope that unraveling the mechanisms involved in the interaction between induced neuronal differentiation of mesenchymal stem cells and PUFA metabolism will help clarify the roles of the exceptionally high proportion of ARA and DHA in neuronal tissue.

## Abbreviations

ACSL: acyl-CoA synthetase long-chain family member;
ARA: arachidonic acid;
bFGF: basic fibroblast growth factor;
BF_3_: boron trifluoride;
BHA: butylated hydroxyanisole;
DAPI: 4,6-diamidino-2-phenylindole;
dbcAMP: dibutyryl cyclic AMP;
DHA: docosahexaenoic acid;
EGF: epidermal growth factor;
EPOX: epoxygenases;
FABP: fatty acid binding protein;
FAME: fatty acid methyl ester;
FCS: fetal calf serum;
GAPDH: glyceraldehyde 3-phosphate dehydrogenase;
IBMX: 3-isobutyl-1-methyl-xanthine;
LOX: lipoxygenases;
MSC: mesenchymal stem cells;
NeuN: anti neuronal nuclei;
NGF: nerve growth factor;
PL: phospholipid;
PUFA: polyunsaturated fatty acids;
PtdCho: phosphatidylcholine;
PtdEtn: phosphatidylethanolamine;
PtdSer: phosphatidylserine;
PtdIns: phosphatidylinositol;
PLA_2_: phospholipase A_2_;
PCA: principal components analysis;
PGI_2_: prostacyclin;
PG: prostaglandin;
PGE_2_: prostaglandin E_2_;
RA: retinoic acid

## Acknowledgments

This work was supported in part by the Public Committee for the Designation of Estate Funds, the Ministry of Justice, Israel (P.G., D.O.), by the Devora Eleonora Kirshman Fund for Research of Parkinson’s Disease, Tel Aviv University and by the Norma and Alan Aufzein chair of Research of Parkinson’s Disease. This work was performed in partial fulfillment of the requirements for a Ph.D. degree of I.K., Sackler Faculty of Medicine, Tel Aviv University, Israel. The authors thank Ms. Aviva Kluska at FMRC, Dr. Shlomo Bulvik at Laniado Medical Center, Natanya, Israel, Dr. Metsada Pasmanik-Chor and Dr. Varda Oron-Karni, Tel-Aviv University Bioinformatics Unit, Tel Aviv.

---

<u>Figure 6. Differential expression of PtdCho and PtdSer metabolizing genes.</u> The expression of the genes was determined by real-time PCR, as specified in Materials and Methods (Representative result from 3 independent experiments). Values are expressed as arbitrary units (AU) relative to the expression of GAPDH + SD. *p<0.05.

CCTbeta, CTP:phosphocholine cytydyltransferase beta, PSI & II Synth., PtdSer synthase I&II

**Supplements Table 1:** Phospholipase family

| Enzyme | Accession number | Pv +Diff. vs - Diff. | Fold Change |
| --- | --- | --- | --- |
| Phospholipase A2, group IB (pancreas; PLA2G1B) | NM_000928 | 0.117004 | 1.15329 |
| Phospholipase A2, group IIA (platelets, synovial fluid; PLA2G2A) | NM_000300 | 0.0556868 | 1.20063 |
| Phospholipase A2, group IID (PLA2G2D) | NM_012400 | 0.438031 | 1.07725 |
| Phospholipase A2 group IIE (PLA2G2E) | NM_014589 | 0.0410931 | -1.13234 |
| Phospholipase A2 group IIF (PLA2G2F) | NM_022819 | 0.0754624 | 1.11205 |
| Phospholipase A2 group III (PLA2G3) | NM_015715 | 0.0198896 | 1.16948 |
| Phospholipase A2, group IVA (cytosolic, calcium-dependent; PLA2G4A) | NM_024420 | 0.00253971 | 4.28091 |
| Phospholipase A2, group IVB (cytosolic; PLA2G4B) | NM_005090 | 0.0236319 | 1.15076 |
| Phospholipase A2, group IVC (cytosolic, calcium-independent PLA2G4C) | NM_003706 | 0.00136685 | 1.8867 |
| Phospholipase A2, group IVD (cytosolic; PLA2G4D) | NM_178034 | 0.519855 | 1.07338 |
| Phospholipase A2, group IVE (PLA2G4E) | NM_001080490 | 0.577801 | 1.03791 |
| Phospholipase A2, group IVF (PLA2G4F) | NM_213600 | 0.0130975 | 1.12463 |
| Phospholipase A2, group V (PLA2G5) | NM_000929 | 0.24853 | -1.63913 |
| Phospholipase A2, group VI (cytosolic, calcium-independent (PLA2G6) | NM_003560 | 0.322225 | 1.09351 |
| Phospholipase A2, group VII (platelet activating factor acetylhydrolase; PLA2G7) | NM_005084 | 0.139557 | 1.3275 |
| Phospholipase A2, group X (PLA2G10) | NM_003561 | 0.128626 | 1.17302 |
| Phospholipase A2, group XIIA (PLA2G12A) | NM_030821 | 0.0338763 | -1.23194 |
| Phospholipase A2, group XIIB (PLA2G12B) | NM_032562 | 0.0800661 | 1.05786 |
| Phospholipase B1 (PLB1) | NM_153021 | 0.205506 | 1.09005 |
| Phospholipase C, beta 1 (phosphoinositide-specific; PLCB1) | NM_182734 | 0.000229454 | 3.68169 |
| Phospholipase C, beta 2 (PLCB2) | NM_004573 | 0.227972 | 1.09475 |
| Phospholipase C, beta 3 (phosphatidylinositol-specific; PLCB3) | NM_000932 | 0.0169165 | -1.18872 |
| Phospholipase C, beta 4 (PLCB4) | NM_000933 | 0.0288829 | -2.83947 |
| Phospholipase C, gamma 1 (PLCG1) | NM_002660 | 0.00668083 | 1.40432 |
| Phospholipase C, gamma 2 (phosphatidylinositol-specific; PLCG2) | NM_002661 | 0.0168439 | 1.2383 |
| Phospholipase C, delta 1 (PLCD1) | NM_006225 | 0.413757 | -1.15326 |
| Phospholipase C, delta 3 (PLCD3) | NM_133373 | 0.0333648 | -1.19087 |
| Phospholipase C, delta 4 (PLCD4) | NM_032726 | 0.538663 | 1.16461 |
| Phospholipase C, epsilon 1 (PLCE1) | NM_016341 | 0.00386173 | -5.53743 |
| Phospholipase C, zeta 1 (PLCZ1) | NM_033123 | 0.0425366 | 1.072 |
| Phospholipase C, eta 1 (PLCH1) | NM_014996 | 0.179965 | 1.08134 |
| Phospholipase C, eta 2 (PLCH2) | NM_014638 | 0.158536 | -1.08059 |
| Phospholipase D1, (phosphatidylcholine-specific; PLD1) | NM_002662 | 0.0023179 | 4.65278 |
| Phospholipase D2 (PLD2) | NM_002663 | 0.0174509 | 1.64138 |
| Phospholipase D family, member 3 (PLD3) | NM_001031696 | 0.0353565 | 1.11496 |
| Phospholipase D family, member 4 (PLD4) | NM_138790 | 0.847484 | 1.01576 |
| Phospholipase D family, member 5 (PLD5) | NM_152666 | 0.452104 | 1.07015 |

**Table 2:** Cyclooxygenase pathway

| Enzyme | Process | Accession number | Pv +Diff. vs - Diff. | Fold Change |
| --- | --- | --- | --- | --- |
| Prostaglandin G/H (PGH<sub>2</sub>) synthase 1 [Cyclooxygenase 1 (COX-1)] | ARA → PGG<sub>2</sub>/PGH<sub>2</sub> | NM_000962 | 0.106273 | 1.33669 |
| PGH<sub>2</sub> synthase 2 [COX-2] | ARA → PGG<sub>2</sub>/PGH<sub>2</sub> | NM_000963 | 8.23E-07 | 41.7132 |
| Prostacyclin (PGI<sub>2</sub>) synthase 2 | PGH<sub>2</sub> → PGI<sub>2</sub> | NM_000961 | 8.42E-06 | -9.27937 |
| Prostaglandin E (PGE<sub>2</sub>) synthase | PGH<sub>2</sub> → PGE<sub>2</sub><br>(key enzyme) | NM_004878 | 1.35E-08 | 15.2885 |
| Prostaglandin E (PGE<sub>2</sub>) synthase 2 | PGH<sub>2</sub> → PGE<sub>2</sub> | NM_025072 | 0.143423 | 1.10351 |
| Prostaglandin E (PGE<sub>2</sub>) synthase 3 | PGH<sub>2</sub> → PGE<sub>2</sub> | NM_006601 | 0.0234323 | -1.22191 |
| Prostaglandin D (PGD<sub>2</sub>) synthase (21kDA brain) | PGH<sub>2</sub> → PGD<sub>2</sub> | NM_000954 | 0.0368164 | -1.36101 |
| Prostaglandin D (PGD<sub>2</sub>) synthase hematopoietic | PGH<sub>2</sub> → PGD<sub>2</sub> | NM_014485 | 0.44254 | 1.04544 |
| Prostaglandin F (PGF<sub>2</sub>) synthase | PGH<sub>2</sub> → PGF<sub>2α</sub> | NA | NA | NA |
| Thromboxane A<sub>2</sub> (TXA<sub>2</sub>) synthase 1 | PGH<sub>2</sub> → TXA<sub>2</sub> | NM_001061 | 0.00652478 | 1.48198 |

**Table 3:**
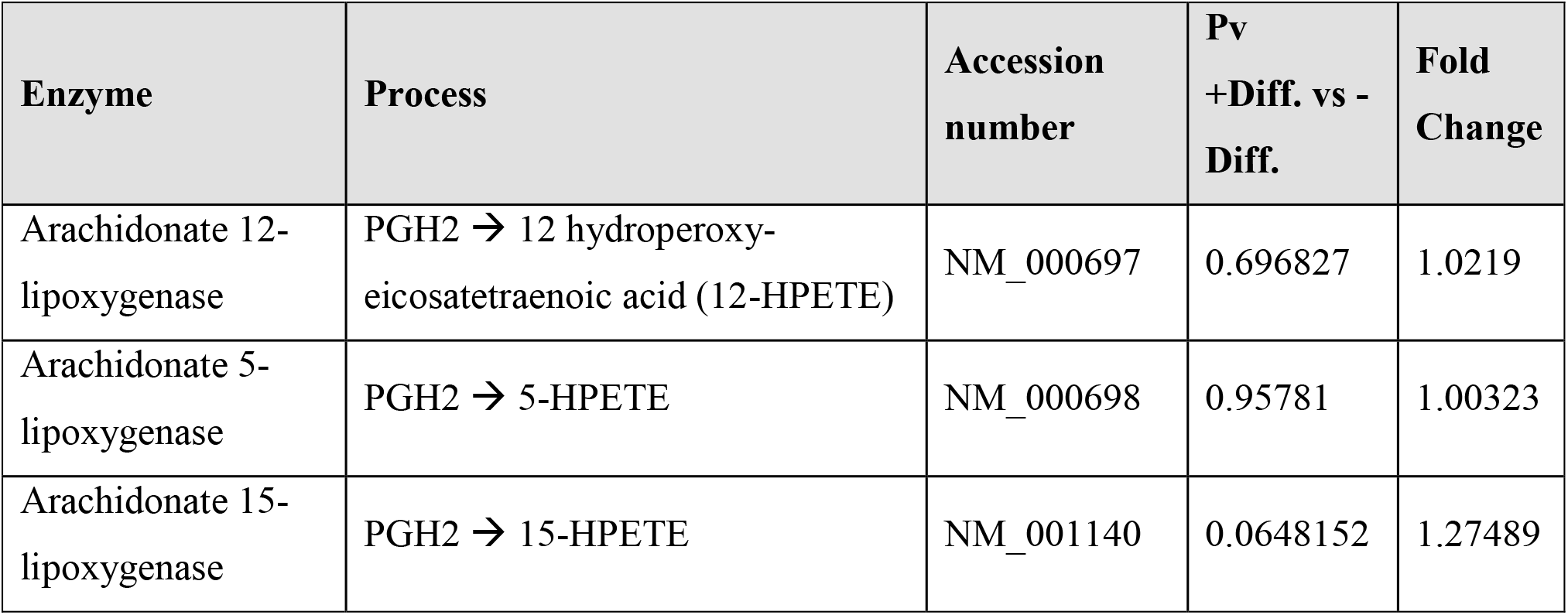
Lipoxygenase pathway

**Table 4:** Cytochrome P450 pathway.

| **Enzyme** | **Process** | **Accession number** | **Pv +Diff. vs - Diff.** | **Fold Change** |
| --- | --- | --- | --- | --- |
| Cytochrome P450 family 51 (CYP51A1) | PGH2 → epoxy-eicosatrienoic acids (EETs) | NM_000786 | 0.000712437 | -1.84349 |

**Table 5:** Fatty acid binding proteins and long chain family member acyl-coA

| **Enzyme** | **Accession number** | **Pv +Diff. vs -Diff.** | **Fold Change** |
| --- | --- | --- | --- |
| Fatty acid binding protein (FABP) 1, liver | NM_001443 | 0.32333 | 1.05729 |
| FABP 2, intestinal | NM_000134 | 0.833212 | 1.01753 |
| FABP 7, brain | NM_001446 | 0.14977 | 1.06952 |
| FABP 3, muscle and heart | NM_004102 | 0.0425585 | -3.16309 |
| FABP 4, adipocyte | NM_001442 | 0.104563 | 1.07656 |
| FABP 5 (psoriasis-associated)(EFABP) | NM_001444 | 0.00000800269 | 10.2446 |
| FABP 6, ileal (gastrotropin) | NM_001445 | 0.0218595 | 1.17291 |
| FABP 7, brain | NM_001446 | 0.14977 | 1.06952 |
| Acyl-CoA synthetase long-chain family member (ACSL) 1 | NM_001995 | 0.000000610651 | 5.77037 |
| ACSL 3 | NM_004457 | 0.00106515 | 2.94836 |
| ACSL 4 | NM_022977 | 0.0000660307 | 11.4308 |
| ACSL 5 | NM_203380 | 0.0369948 | -1.31453 |
| ACSL 6 | NM_001009185 | 0.229006 | 1.01938 |

## Notes

### Competing Interest Statement

The authors have declared no competing interest.

